# Endothelial Protein C Receptor Could Contribute to Experimental Malaria-Associated Acute Respiratory Distress Syndrome

**DOI:** 10.1101/348318

**Authors:** Luana dos Santos Ortolan, Michelle Klein Sercundes, Gabriel Candido Moura, Thatyane de Castro Quirino, Daniela Debone, Douglas de Sousa Costa, Oscar Murillo, Claudio Romero Farias Marinho, Sabrina Epiphanio

## Abstract

The severity of *Plasmodium falciparum* malaria is associated with parasite cytoadherence, but there is limited knowledge about the effect of parasite cytoadherence in malaria-associated acute respiratory distress syndrome (ARDS). Our objective was to evaluate the cytoadherence of infected red blood cells (iRBCs) in a murine model of ARDS and to appraise a potential function of endothelial protein C receptor (EPCR) in ARDS pathogenesis. DBA/2 mice infected with *P. berghei* ANKA were classified as ARDS- or hyperparasitemia (HP)-developing mice according to respiratory parameters and parasitemia. Lungs, blood and bronchoalveolar lavage were collected for gene expression or protein analyses. Primary cultures of microvascular lung endothelial cells from DBA/2 mice were analyzed for iRBC interactions. Lungs from ARDS-developing mice showed evidence of iRBC accumulation along with an increase in EPCR and TNF concentrations. Furthermore, TNF increased iRBC adherence *in vitro*. Dexamethasone-treated infected mice showed low levels of TNF and EPCR mRNA expression and, finally, decreased vascular permeability, thus protecting mice from ARDS. In conclusion, we identified that increased iRBC cytoadherence in the lungs underlies malaria-associated ARDS in DBA/2-infected mice and that inflammation increased cytoadherence capacity, suggesting a participation of EPCR and a conceivable target for drug development.

## INTRODUCTION

Malaria infection by *Plasmodium falciparum* is responsible for the largest number of severe and fatal diseases in the tropics [1,2]. The main complications of *P. falciparum* infection include cerebral malaria, pulmonary complications, acute renal failure, severe anemia, bleeding and placental malaria [3]. An important aspect of the pathogenesis of severe malaria results from the ability of infected red blood cells (iRBCs) to adhere to the microvasculature. This interaction between iRBCs and the endothelium can cause blocking of blood flow and/or a local inflammatory response [3–5]. Furthermore, these adhesions promote the disappearance of asexual forms of the parasite in the peripheral circulation, thus preventing them from being destroyed in the spleen [3,5,6]. Pulmonary complications caused by severe malaria include acute respiratory distress syndrome (ARDS), which has been associated with not only severe malaria but also different diseases [7–9]. Although malaria-associated ARDS often causes a high mortality rate, not so much research has been performed. Murine models have been used to study malaria-associated ARDS [10,11], and DBA/2 mice infected with *P. berghei* ANKA (PbA) develop ARDS and die between the 7^th^ to 12^th^ days post-infection (dpi) with pleural effusion, edema and inflammatory infiltration in the lungs but without signals of cerebral malaria. In contrast, DBA/2 mice that died after the 13^th^ dpi exhibited pale lungs, no pleural effusion and high levels of parasitemia. The cause of death was attributed to hyperparasitemia (HP) and consequent anemia. In addition, we established predictive criteria to distinguish which mice would die from ARDS on the 7^th^ dpi using respiratory and parasitemia data [12]. Using this model, we found that on average 50% of the mice died from ARDS and 50% died from HP. We found that recruitment of neutrophils and vascular endothelial growth factor (VEGF) is essential to the pathogenesis of malaria-associated ARDS [13,14] and that the induction of heme oxygenase-1 (HO-1) has a protective effect against the development of ARDS in mice [15]. Epiphanio *et al* showed sFLT1 (soluble form of VEGF receptor, known to neutralize excess VEGF in circulation) - expressing adenoviruses treated mice protected against ARDS which was correlated with significant decrease in VEGF levels in circulation, even as carbon monoxide administration by inhalation suppresses the onset of this syndrome [13]. In addition, it was demonstrated that hemin (inducer of HO-1) treated mice had also VEGF levels reduced and were protected against ARDS, defending the alveolar capillary barrier in vitro [15]. Although VEGF is important in PbA-infection of DBA/2 mice, this is not a universal effect, since neutralization of VEGF receptor-2 did not decrease ARDS pathology in *P. berghei* NK65-infected C57BL/6 mice, another well-defined and accepted model of ARDS in malaria [16].

Additionally, it has already been demonstrated that PbA-iRBCs adhere to MVECS (microvascular lung endothelial cells from CBA/Ca mice) and that TNF-stimulated cells express more ICAM-1 and VCAM than control cells [17]. In the *P. chabaudi* model, infected mice in the absence of ICAM-1 showed less anemia and weight loss, reduced parasite accumulation in both the spleen and liver and higher peripheral blood parasitemia during acute stage malaria, which presented the possible role of ICAM-1 in adhesion and pathogenesis of *P. chabaudi* [18]. Endothelial receptors have been studied to understand iRBC adherence in the microvasculature of different organs in severe malaria, and some of these receptors are well established such as ICAM-1 in cerebral malaria and CSA in placental malaria [19]. However, both the mechanism and consequences of PbA-iRBC adhesion in DBA/2 lung endothelial cells remain unknown. In 2013, endothelial protein C receptor (EPCR) was shown to be a new receptor for *Plasmodium falciparum* erythrocyte protein 1 (PfEMP1) in *P. falciparum-*induced severe malaria [20], which may result in a new branch of research on severe malaria. As its primary function, EPCR binds with activated protein C (APC) and cleaves protease-activated receptor 1 (PAR-1) in a specific Ras-related C3 botulinum toxin substrate 1 (RAC1) pathway, which inhibits the activation of nuclear factor-κB and provides barrier protection [21]. The EPCR facilitates the activation of protein C (PC) by the thrombin-thrombomodulin complex, promoting cytoprotective effects in vessels and tissue protection in the brain, lungs, kidneys, and liver [22]. Additionally, APC provides neuroprotective effects such as anti-inflammatory and anti-apoptotic effects and protection of the blood-brain barrier, kidneys, and lungs and thus may be directly relevant to the complications associated with severe malaria [23].

Some authors proposed that in malaria, *P. falciparum*-infected erythrocytes prompt a decrease in EPCR levels, and, consequently, APC production is disabled, resulting in enhanced coagulation and inducing proinflammatory factors and endothelial dysfunction via PAR-1 [24]. Indeed, *in vitro* studies show that purified amino-terminal cysteine-rich interdomain region (CIDRα1), especially that from domain cassette 8 (DC8), member of the group containing PfEMP1, interferes with protein C binding to EPCR, resulting in acquired functional PC systemic deficiency [20,23,25]. However, it has unexpectedly been shown that iRBCs expressing DC13 and the HB3var03 or IT4var07 variants of PfEMP1 do not bind to the EPCR of brain endothelial cells *in vitro*. On the other hand, it has been shown that the DC8 variant IT4var19 may bind to the EPCR, but this interaction was inhibited when human serum or plasma was added to the assay [25]. Therefore, the disagreement concerning PfEMP1-EPCR interactions indicates the need for further studies to understand the outcome of severe malaria. Notwithstanding, there is no evidence to date of *P. berghei* ANKA binding to EPCR, and the effects of EPCR on ARDS pathogenesis must be elucidated. Our study, therefore, was developed in a murine model that mimics various types of human ARDS [12] and in primary culture of microvascular lung endothelial cells from DBA/2 mice (PMLEC) not only to clarify the adhesion of infected erythrocytes to murine lung microvascular endothelial cells but also to understand the relevant aspects of EPCR modulation by the immune response, which can bring important contributions to understanding malaria-associated ARDS.

## MATERIALS AND METHODS

### Experimental outline

DBA/2 mice were infected with 10^6^ infected red blood cells (iRBC) with *Plasmodium berghei* ANKA (PbA) and classified as ARDS-developing or HP-developing mice before death according to previously described criteria [12]. Briefly, we used two groups of infected mice: the survival group (infected control) and the euthanized group, in which the mice were euthanized on the 7^th^ day postinfection (10–12 mice per group). By using respiratory patterns (enhanced pause and respiratory frequency) and the degree of parasitemia as predictive criteria, we established cut-off values using receiver operating characteristic (ROC) curves for these parameters measured on the 7^th^ dpi based on data from mice whose cause of death was known (survival group). In the survival group, for mice showing pleural effusion or red and congested lungs at necropsy, the cause of death was designated as ARDS. For mice without pleural effusion that died after 13 days postinfection with pale lungs and high levels of parasitemia, the cause of death was designated as hyperparasitemia (HP). Afterward, we retrospectively diagnosed the euthanized mice as suffering from ARDS or HP by comparing their respiratory patterns and parasitemia measured on the 7^th^ dpi with the cut-off values from the survival group at the end of each experiment (20^th^ dpi) [14,15]. All lung tissue, blood, and bronchoalveolar lavage (BAL) samples were collected from mice on the 7^th^ day postinfection (dpi) always after perfusion of the right ventricle with 20 mL of 1× PBS until the lungs remained clear.

### Mice, parasites and euthanasia

Male DBA/2 mice between 6-10 weeks old (purchased from the Department of Parasitology, University of São Paulo, Brazil) were infected with 1 ×10^6^ *P. berghei* ANKA (clone 1.49L) iRBCs kindly provided by the laboratory of Dr. Maria Mota from the Institute of Molecular Medicine (IMM) in Portugal. Parasitemia and mortality were monitored daily. Parasitemia was determined by Giemsa staining and expressed as the percentage of infected red blood cells. The euthanasia of mice was performed using ketamine (150 mg/kg)/xylazine (15 mg/kg).

### Determination of Respiratory Pattern

Respiratory patterns (respiratory frequency [RF], tidal volume [TV] and enhanced pause [Penh]) were monitored on the 7^th^ dpi by an unrestrained whole-body plethysmography chamber (WBP, Buxco Electronics, USA) for 10 minutes (basal level) according to previously described methods [12].

### Histopathological analyses and hemozoin count

Lung tissue fragments were fixed with 10% buffered formalin for 24 hours and kept in 70% ethanol until being embedded in paraffin, and 4–5 μm sections were stained with hematoxylin-eosin (H&E). To determine the hemozoin (Hz)-containing area, lungs from euthanized mice (ARDS-developing or HP-developing mice according to the predictive model on the 7^th^ day postinfection [12]) were stained with H&E, and 10 images were captured from each tissue sample with polarized light (400× magnification) using a Zeiss color camera (CAM Axio HRc) connected to a Zeiss light microscope (Axio Imager.M2). The corresponding percentage of Hz in each image was identified using the program ImageJ, in which the areas containing Hz were distinguished by brightness adjustment. The area containing Hz was normalized by the total area in each image.

### Quantification of gene expression

Quantitative RT-PCR was performed for the relative quantification of gene expression from the lungs of noninfected and infected mice. RNA extraction was performed according to the “Animal Cell I” protocol from a RNeasy Mini kit (Qiagen, USA). cDNA synthesis was performed with a 1 μg RNA sample using the First Strand cDNA Synthesis kit RT-PCR (Roche, USA) according to the manufacturer’s instructions. Finally, the PCR sample to test gene expression was prepared with SYBR Green PCR Master Mix (Applied Biosystems, USA), and the relative quantification 2 ^(-ΔΔCT)^ method was used as described before (51). The qRT-PCR reactions were performed in the ABI 7500 Fast instrument (Applied Biosystems, USA) using the following oligonucleotides: PbA 18S (forward 5’-agcattaaataaagcgaatacatccttac-3’; reverse 5’-ggagattggttttgacgtttatgtg-3’); HPRT (forward 5’-aagcttgctggtgaaaagga-3’; reverse 5’-ttgcgctcatcttaggcttt-3’); ICAM-1 (forward 5’-cgaaggtggttcttctgagc-3’; reverse 5’-gtctgctgagscccctcttg-3’); VCAM (forward 5’-agtccgttctgaccatggag-3’; reverse 5’-tgtctggagccaaacacttg-3’); EPCR (forward 5’-gacgaagtttctgccgctac-3’; reverse 5’-ctggaggatggtgacgtttt-3’).

### Parasite localization by bioluminescence

DBA/2 mice were infected with luciferase-expressing *P. berghei* parasite (parasite line 354cl4) kindly provided by the laboratory of Dr. Maria Motta from the Institute of Molecular Medicine (IMM) in Portugal. On the 7^th^ day postinfection, 150 µl of luciferin (2.25 mg) (VivoGlo ™ Luciferin, In Vivo Grid, Catalog #: P1041, Promega) was injected intraperitoneally, allowing the emission of parasite luminescence. Then, the localization of the parasite was analyzed with an IVIS Spectrum system (PerkinElmer). For this purpose, mice were sedated with isoflurane for pictures (approximately 6 minutes after the injection of luciferin). Later, they were euthanized and perfused in the right ventricle with 20 ml of 1× PBS. After perfusion, a new image of the mice was captured, and the organs were collected and placed in sterile Petri dishes to observe the bioluminescence of each tissue.

### Cytokines quantification

The quantities of TNF, IL-6, and IL-33 in serum and culture supernatants were determined by Mouse ELISA Ready-SET-Go! ^®^ commercial kits from eBioscience (San Diego, CA, USA) specific for TNF (ref 88-7324-88), IL-6 (ref 88-7064-88) and IL-33 (ref 88-7333-88) according to the manufacturer’s protocols.

### Isolation of the primary microvascular lung endothelial cells

Primary microvascular lung endothelial cells (PMLECs) were obtained from naive DBA/2 mice as described before (15,52). Briefly, after euthanasia, the body of the animal was disinfected with iodine/alcohol. Then, the mice had all of their blood removed by cutting the carotid artery. In a laminar flow chamber, the lung tissue was cut into fragments of approximately 1 mm^2^ and distributed among 6-well polystyrene plates in low glucose DMEM (Invitrogen)-supplemented culture medium [20% heat inactivated FBS, 40 μg/ml gentamicin (Invitrogen-15710064) and 1:100 antibiotic/antimycotic (Gibco-25200)] at 37°C and 5% CO2. After 72 hours, the tissue fragments were removed, and 50% of the medium was replaced. After 7 days of incubation, the cells were removed with trypsin 0.25% EDTA (Gibco) and replaced in a 75 cm^2^ culture flask. The trypsinization procedure was repeated every 5 to 7 days. Finally, the cells were cultured for 15 to 20 days (3^rd^ and 5^th^ passage) until being used in the trials. The purity of isolated DBA/2-PMLECs was characterized by immunofluorescence with lectin from *Ulex europaeus* (Sigma-Aldrich, USA– l9006) and the anti-VWF (Santa Cruz Biotechnology, USA – sc14014), anti-CD31 (Abcam, UK – ab28364), anti-ACE (Abcam, UK – ab85955), anti-CD62E (Abcam, UK - ab18981), anti-eNOS (Abcam, UK – ab87750) and anti-VE-cadherin (Abcam, UK – ab205336) antibodies. All of the cells were stained with all of the markers, indicating high purity (Figure S1).

### *Plasmodium* synchronization and enrichment of parasitized erythrocytes

To obtain mature forms of *P. berghei* ANKA, iRBCs were synchronized as described previously (53). Briefly, iRBCs were collected from infected mice exhibiting 10 to 20% parasitemia through cardiac puncture and transferred to RPMI 1640 culture medium (Gibco, Thermo Fisher Scientific, USA) supplemented with 25% fetal bovine serum (FBS). The iRBCs were subsequently maintained *in vitro* at 37°C for 14 hours in an atmosphere containing 5% CO_2_, 85% N_2_, and 10% O_2_. The parasitized erythrocytes were then enriched using a magnetic separation column (Miltenyi Biotec, USA) to generate cell populations consisting of approximately 95% iRBCs, as assessed by thick blood smears. Lysate containing *P. berghei* was obtained from iRBCs subjected to several freeze-thaw cycles.

### Peritoneal Macrophages

Peritoneal macrophages (Mφ) were collected from noninfected DBA/2 mice euthanized with halothane, the skin was then removed and 5 ml of 1× sterile cold PBS (at 4°C) was inserted into the peritoneum. The liquid was then removed, aspirated with a 24-gauge needle, and centrifuged at 1000 rpm for 5 minutes at 4°C. The supernatant was discarded, and the cells were resuspended in supplemented DMEM medium [20% heat-inactivated FBS, 40 μg/ml gentamicin (Invitrogen-15710064) and 1:100 antibiotic and antimycotic (Gibco-25200)]. Cells were counted in a Neubauer chamber and arranged in Transwell culture plates pretreated with 0.2% gelatin in 1× PBS (gelatin from bovine skin, G9391, Sigma-Aldrich).

### Transwell assay with endothelial cells and peritoneal macrophages

Sterile 13 mm diameter coverslips (Knittel) were placed in 24-well culture plates. The coverslips were treated with 0.2% gelatin (gelatin from bovine skin - Sigma-Aldrich) diluted in 1× PBS. PMLECs were seeded in the culture plate (7 × 10^4^ cells/well) in 400 μl of complete DMEM medium, and 1× 10^5^ peritoneal macrophages (Mφ) were seeded in the Transwell membrane (6.5 mm membrane diameter, 0.4 μm membrane pore, Corning, Costar 3470) without contacting PMLECs on the bottom of the 24-well plate. They only shared the supernatant through the Transwell membrane. After 24 hours, 2.5 × 10^6^ mature iRBCs or RBCs were added over the Mφ and coincubated for 24 hours. After this time, the supernatant between the Mφ and endothelial cells was collected for the quantification of cytokines by ELISA. Subsequently, endothelial cells were coincubated with 1.75 × 10^6^ PbA-iRBCs for the adhesion assay.

### *Plasmodium berghei* ANKA adhesion assay under static conditions

DBA/2-PMLECs were plated in a Lab-Tek chamber slide system made of Permanox containing 8 wells (3 ×10^4^ cells/well) (Thermo Fisher Scientific - 177 455). Afterwards, TNF (50 ng/ml) was added to the cells and incubated for 24, 48 or 72 hours. After this stimulation, synchronized PbA-iRBCs were added for 1 hour to the culture at a ratio of 25 iRBCs/PMLEC at 37°C and 5% CO_2_. After the incubation period, the culture medium and the removable chamber were removed, and the slides were dipped in preheated DMEM medium without FBS to eliminate unbound PbA-iRBCs. The slides were then fixed with methanol, stained with Giemsa and observed under an optical microscope immersion objective (1000×). The counting pattern used was the number of adhered PbA-iRBCs for every 100 DBA/2-PMLECs.

### Parasite adhesion assay in flowing conditions

PMLECs between the 3^rd^ and 5^th^ passages were seeded (8 × 10^4^/well) in 2-well Permanox chamber slides from Lab-tek (Thermo Scientific, Nunc). The cells were incubated at 37°C and 5% CO_2_ for 24 hours. After this time, TNF was added for 24 hours or not added (for the control group). The stimulus was removed, and mature PbA-iRBCs (25 per cell) in DMEM supplemented with 20% FBS were added and allowed to interact with the endothelial cells for 1 hour at 37°C in 5% CO_2_. Subsequently, the wells were detached from the polystyrene slides, and the slides were added to a flow system composed of a chamber (cell adhesion flow chamber, Immunetics) that kept the slide adhered through the formation of a vacuum (coupled by a vacuum pump), a syringe pump (Insight Inc.), an inverted microscope (Zeiss Vert. A1) connected to a camera (Axio Cam ERc 5s, Zeiss) and a computer with an image capture system (Zen 2011 program, AxioVision Rel 4.8.2 SP2). The chamber was initially filled with DMEM medium, and then an initial photo with the parasites adhered to the PMLEC-DBA/2 cells was taken. Then, a medium continuous stream was run through the chamber and maintained at a flow rate of 2 ml/hour with a pump syringe (Insight Inc.). Five images, taken every 3 minutes, of each well were captured while maintaining the same field as the initial image to determine the erythrocyte binding efficiency. Image processing and analysis were performed using ImageJ (version 1.46 r). The images were opened individually in ImageJ and analyzed through the “cell counter” plugin, and all PbA-iRBC images were selected on each image. After counting all the images, the total number of PbA-iRBCs that remained in the final image (taken after 15 minutes) was recorded. The experiment was repeated with two independent cultures, each with two technical replicates. Image processing and analysis were performed in ImageJ.

### Treatment with dexamethasone

A total of 200 μl (80 mg/kg) of dexamethasone (8 mg/ml Decadronal, Aché, Brazil) was taken directly from the bottle and injected intraperitoneally (ip) on days 5 and 6 postinfection. The dexamethasone dose was chosen according to the therapeutic dose observed by Van den Steen in 2010 [29]. Control mice received 200 μl of 1× PBS ip on the same days.

### Lung permeability and edema quantification

To investigate lung permeability, noninfected or PbA-infected mice (dexamethasone-treated or not) on the 7^th^ dpi were injected intravenously with 0.2 mL of 1% Evans Blue (Sigma-Aldrich). The mice were euthanized 45 minutes later, and the lungs were weighed immediately and placed in 2 mL of formamide (Merck) for 48 hours at 37°C [12]. The absorbance of the formamide was then measured at 620 nm and 740 nm. The amount of Evans Blue staining per gram of lung tissue was calculated from a standard curve.

### Measure of primary microvascular lung endothelial cell permeability

The increased lung vascular permeability was analyzed in DBA/2-PMLECs plated on permeable membrane inserts with 0.4 µM pores (Transwell Corning) pretreated with gelatin 0.2% in 1× PBS (gelatin from bovine skin, G9391, Sigma-Aldrich), coupled in 24-well polystyrene plates at a concentration of 2.2 × 10^4^ cells per insert and maintained in DMEM culture at 37°C as previously described (15). After 96 hours, when the cells reached confluency, PbA lysate was applied for 1 hour after incubation with dexamethasone (500 ng/ml for 24 h) or solely with 20% FBS-supplemented DMEM culture medium. The culture medium was subsequently replaced by Hank’s balanced salt solution, and in the upper compartment of each insert in contact with the cells, 200 µl of Evans Blue was incubated at a 2 mg/mL concentration at 37°C. After 30 minutes, the liquid from the lower compartment was collected and analyzed in a spectrophotometer at a wavelength of 650 nm (NanoDrop 2000, Thermo Scientific). Finally, the concentration of Evans Blue was determined from a standard curve (0.2 mg/mL to 0.0031 mg/mL) as previously described [30].

### Actin microfilament identification by immunofluorescent and morphometric analysis of the opening of interendothelial junctions

To analyze the area of opening of interendothelial junctions (OIJ), PMLECs were plated in 24-well plates (7×10^4^ cells/well), adhered to gelatin on glass coverslips, and maintained at 37°C and 5% CO_2_. The cells were coincubated with either iRBCs or RBCs for 1 hour after incubation with dexamethasone (24 hours) or DMEM culture medium supplemented with 20% FBS, and incubations were done in triplicate. Subsequently, the cells were fixed with 3.7% formaldehyde, permeabilized with acetone at −20°C, and blocked with bovine serum albumin solution (1% BSA). Actin was marked with Texas Red-phalloidin (T7471, Life Technologies) for 20 minutes. The cell nuclei were marked with Hoechst stain (H33342, Life Technologies). Each slide with fully confluent cells was chosen randomly, and ten to twenty pictures were taken and scanned in a “zig-zag” pattern from top to bottom. The images were acquired with an Axio Imager M2 microscope (Zeiss) using Axiocam HRc (Zeiss) and Axio Vision software, version 4.9.1.0. The total OIJ area was measured in each picture using ImageJ software (version 1.52a - NIH, USA). To calculate the OIJ, the “measure” tool and the “wand (tracing)” tool in ImageJ were used. Each image was analyzed individually, and all of the OIJs were circled with the “wand (tracing)” tool, from which the area of each space was calculated. Analysis of the differences between the openings of the interendothelial junctions following different stimuli was performed using GraphPad Prism 8^®^ software.

### TNF blockage

TNF was blocked in DBA-PMLECs using mouse TNF neutralizing (D2H4) rabbit mAb #11969 (Cell Signaling^®^) at a concentration of 1 µg/ml for 24 hours in PMLECs.

### Soluble EPCR quantification

Soluble endothelial protein C receptor (sEPCR) was measured with an ELISA kit (Elabscience^®^, E-EL-M1073) according to the manufacturer’s instructions.

### EPCR Western Blot

Fresh frozen mouse lung tissues collected on the 7^th^ dpi were sonicated and homogenized at 4°C using Radio-Immunoprecipitation Assay (RIPA) buffer composed of 50 mM Tris-HCl pH 8.0, 150 mM NaCl, 0.5% sodium deoxycholate, 0.2% sodium dodecyl sulfate (SDS), 1 mM sodium orthovanadate, 1 mM NaF, and a protease inhibitor tablet. The total protein concentration was determined using a SpectraMax^®^ Plus 384 (Molecular Devices) spectrophotometer with a 562 nm filter using the Pierce™ BCA Protein Assay kit (Thermo Scientific) according to the manufacturer’s instructions. Then, 30 μg of protein from each sample was electrophoretically separated in a 10% polyacrylamide mini-gel (Bio-Rad) by SDS-PAGE at 100 V for 2 hours in a Mini-PROTEAN^®^ Tetra cell apparatus (Bio-Rad). The Kaleidoscope commercial standard Precision Plus Protein Standard (Bio-Rad) was used. Subsequently, proteins separated on the gel were transferred to PVDF (polyvinylidene difluoride, 0.2 μm, Bio-Rad) membranes in transfer buffer (0.1 M Tris, 20% methanol and MilliQ water) overnight at 4°C in a wet transfer apparatus at 70 mA constant voltage (Bio-Rad). Detection of the chemiluminescence immunochemical reaction was performed on a ChemiDoc ™ XRS + Molecular Imager^®^ (Bio-Rad) transilluminator with the ImageLab ™ program (Bio-Rad). Blocking of nonspecific sites was carried out after the transfer in Tris-buffered saline and Tween-20 buffer (0.05 M Tris, 0.1 M NaCl, pH 7.3, and 0.1% Tween-20, Synth) containing 3% BSA for 2 hours while stirring at room temperature (RT). The membrane was then incubated with the primary anti-EPCR antibody (#151403, Abcam) at 1:1,000 while stirring overnight at 4°C. After washing in TBS-T, the membrane was incubated with the peroxidase-conjugated rabbit anti-rabbit IgG (HRP) secondary antibody (#AP307P, Millipore) at 1:6,000 in TBS-T for 1 hour (RT). Bands corresponding to EPCR and the housekeeping protein *β*-actin (mouse IgG #NB600-501, Novus, at 1:40,000) were detected by the chemiluminescence method (Clarity Western ECL Bio-Rad) and finally densitometrically measured by ImageJ 1.6.0 software.

### Bronchoalveolar lavage and VEGF quantification

On the 7^th^ dpi, infected mice (untreated or treated with dexamethasone) were anesthetized, and having bronchoalveolar lavage (BAL) collected. For that, the trachea was exposed and cannulated, and the lungs were washed once with 1.0 ml of 1× PBS. An ELISA kit (R&D Systems, USA) was used to quantify VEGF levels in BAL and in culture supernatant according to the manufacturer’s instructions.

### Transfection of EPCR by siRNA

To transfect EPCR by interference RNA assay, Lipofectamine® RNAiMAX reagent (Invitrogen by Life Technologies) and 3 different predesigned and validated oligo silencers (Ambion by Life Technologies) were used according to the manufacturer’s protocol. Additionally, a Select Negative Control silencer [(C-) (Ambion 4390843)] and a select GAPDH positive control silencer [(C+) (Ambion 4390849) were employed as control reactions.

siRNA1: forward: 5’ CAACCGGACUCGGUAUGAATT 3’;

reverse: 5’ UUCAUACCGAGUCCGGUUGta3’

siRNA2: forward: 5’ ACGCAAAACAUGAAAGGGATT 3’;

reverse: 5’ UCCCUUUCAUGUUUUGCGUGG 3’

siRNA3: forward: 5’ CGCCCUUUGUAACUCCGAUTT 3’;

reverse: 5’ AUCGGAGUUACAAAGGGCGCA3’

### Immunohistochemistry of ICAM-1 and VCAM in lungs

To perform immunohistochemistry of ICAM-1 and VCAM, slides containing the paraffin tissue sections were placed in an incubator at 60°C for 20 minutes to melt the paraffin. Then, they were incubated in xylene twice for 15□min at 60°C and then in absolute ethanol, 95% alcohol, 70% alcohol, distilled water, and finally 1× PBS, pH 7.2 to 7.4. For antigen retrieval, the slides were incubated in sodium citrate buffer, pH 6, for 45 minutes at 95°C. Endogenous peroxidase was blocked with 3% hydrogen peroxide for 15 minutes twice at room temperature while protected from light. The tissue was probed with rabbit polyclonal to ICAM-1 (1:400) antibody (Abcam, ab124759) and rabbit monoclonal to VCAM (1:600) antibody (Abcam ab134047) overnight at 4°C, and then the REVEAL mouse/rabbit kit (Spring, Code SPD-015) was used in accordance with the manufacturer’s instructions. The quantification of ICAM-1 and VCAM in lung tissue was performed by calculating the marked area normalized to the total tissue section. The calculation was performed in ImageJ (version 1.50b) software using the IHC toolbox plugin [31].

### Statistical Analysis

The statistical analyses were performed using GraphPad Prism^®^ 5.0 software for analysis and graphing. The data were analyzed for normality by the Kolmogorov-Smirnov test or the Shapiro-Wilk normality test and for variance with the Bartlett test. Nonparametric variables for two groups were compared by using the Mann-Whitney test. For analysis of three groups, we used the Kruskal-Wallis test followed by the Dunn’s post hoc test. To compare parametrical variables for two groups, a t-test was employed, and for three or more groups, one-way ANOVA followed by the Bonferroni posttest was used. For the survival curves, log-rank and Gehan-Breslow Wilcoxon tests were applied. The differences between the groups were considered significant when p≤0.05 (5%). To establish a cut-off from the data, ROC curves were generated by using the results of the control group obtained in MedCalc version 8.2.1.0.

### Ethics Statement

All experiments were performed in accordance with the ethical guidelines for experiments with mice, and the protocols were approved by the Animal Health Committee of the Biomedical Sciences Institute of the University of São Paulo (CEUA n^º^ 24, page 16, book 03). The guidelines for animal use and care were based on the standards established by the National Council for Control of Animal Experimentation (CONCEA: Conselho Nacional de Controle de Experimentação Animal) and Brazilian Federal Law n^º^ 11.794.

## RESULTS

### ARDS-developing mice show higher levels of pulmonary parasite than HP- developing mice

In our previous study, it was shown that a proportion of DBA/2 mice infected with PbA-iRBC developed singular characteristics of ARDS and died between the 7^th^ and 12^th^ dpi [12–15]. To investigate whether adherence is essential in the evolution of this phenotype, we observed that luciferase-expressing *P. berghei* parasites were distributed in the peripheral blood and tissues of DBA/2 mice. However, when they were perfused with 1× PBS, the bioluminescence (luciferase/luciferin) signal remained concentrated in the spleen, lungs and slightly in the liver especially in ARDS-developing mice (Figure 1 a-c), corroborating our published data where we showed more bioluminescence signal in ARDS mice compared to HP [14]. We demonstrated that ARDS-developing mice showed higher levels of 18S subunit PbA rRNA expression (Figure 1d) and higher hemozoin concentrations in the lungs compared to those of HP-developing mice on the 7^th^ dpi (Figure 1 e-g). In addition, we analyzed histological lung sections of mice that died with ARDS and found several iRBCs in close contact with endothelial cells (Figure 1h). These results taken together indicated that ARDS-developing mice accumulated a considerable amount of iRBCs in the lungs, suggesting their essential participation in ARDS pathogenesis.

**Figure 1:**
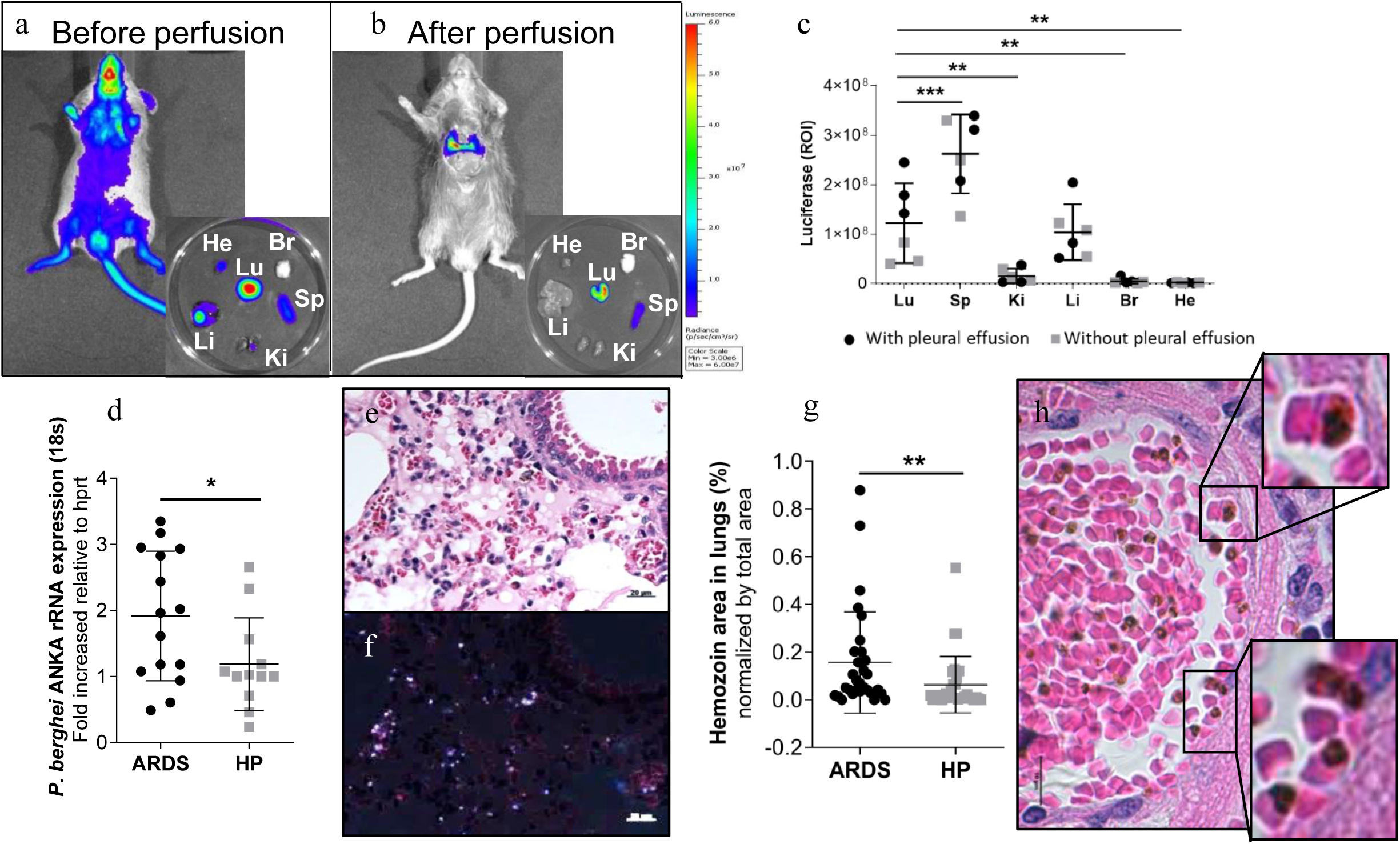
ARDS-developing mice have higher levels of pulmonary parasites than HP-developing mice. (a and b) The distribution of infected red blood cells (iRBCs) was analyzed in DBA/2 mice infected with luciferase-expressing *P. berghei* parasite (PbA-luciferase) on the 7^th^ day postinfection (dpi) and analyzed *in vivo* by IVIS imaging after luciferin. (c) The organs were individually analyzed after perfusion and results were plotted with black circles (mice with pleural effusion) and with gray squares (mice without pleural effusion); one-way ANOVA (**p<0.01; ***p<0.001). (d) Perfused lungs from ARDS- and HP-developing mice were collected on the 7^th^ dpi and analyzed by rRNA gene expression of *P. berghei* ANKA (18S subunit) using the 2^-ΔΔCT^ method. Representative image of a perfused lung collected from an ARDS-developing mouse under (e) normal and polarized lights (f) showing the hemozoin pigments, on the 7^th^ dpi. (g) Quantification of the hemozoin area analyzed by ImageJ (magnification: 400×, scale bar: 20 μm). Bars represent the average ± SD (*p<0.05; **p< 0.01). (d) Unpaired t-test and (g) Mann-Whitney test. (h) Lung histological analyses showing iRBCs in close contact with endothelial cells (inserts) in an infected mouse that died by ARDS (magnification: 630X, scale bar: 10 μm). ARDS: acute respiratory distress syndrome; HP: hyperparasitemia; Lu: lung; Sp: spleen; Ki: kidney; Li: liver; Br: brain; He: heart.

### TNF increases *P. berghei* adherence in the ARDS experimental model

ARDS-developing mice on the 7^th^ dpi displayed higher TNF levels in serum compared to those of HP-developing mice (Figure 2a), suggesting that this inflammatory cytokine may be critical to ARDS development. We further investigated whether iRBCs could contribute to TNF release by endothelial cells. First, we demonstrated that primary microvascular lung endothelial cells (PMLECs) stimulated with iRBCs can directly contribute to TNF production (Figure 2b). Then, we examined the influence of TNF on the adhesion of iRBCs to PMLECs using static and flow conditions. TNF-stimulated cells increased the capacity of iRBCs to adhere to PMLECs after 24 and 72 hours in the static assay (Figure 2 c-e). In the flow adherence assay, which mimics physiologic conditions, the iRBCs had more cytoadherence with TNF than without TNF stimulation (Figure 2 f-h). Additionally, peritoneal macrophages (Mφ), collected from noninfected DBA/2 mice, were seeded in Transwell membranes, and then PMLECs were seeded on the bottom of a 24-well plate with no contact with the Mφ being stimulated or not by red blood cells (RBCs) or iRBCs (Figure 2i-k). Mφ stimulated by iRBCs produced more TNF than Mφ without contact with iRBCs (Figure 2j). Additionally, PMLECs that were in indirect contact with Mφ stimulated with iRBCs demonstrated more adherence to iRBCs than PMLECs with no previous contact with Mφ (Figure 2k). Finally, to evaluate the contribution of TNF to the adhesion of iRBCs to PMLECs, the cells were blocked with TNF antibody, and as a result, the blockage reduced the capacity of iRBCs to adhere to PMLECs (Figure 2l). All these data show that ARDS-developing mice express more TNF, which contributes to iRBC adherence in endothelial cells than in HP-developing mice.

**Figure 2:**
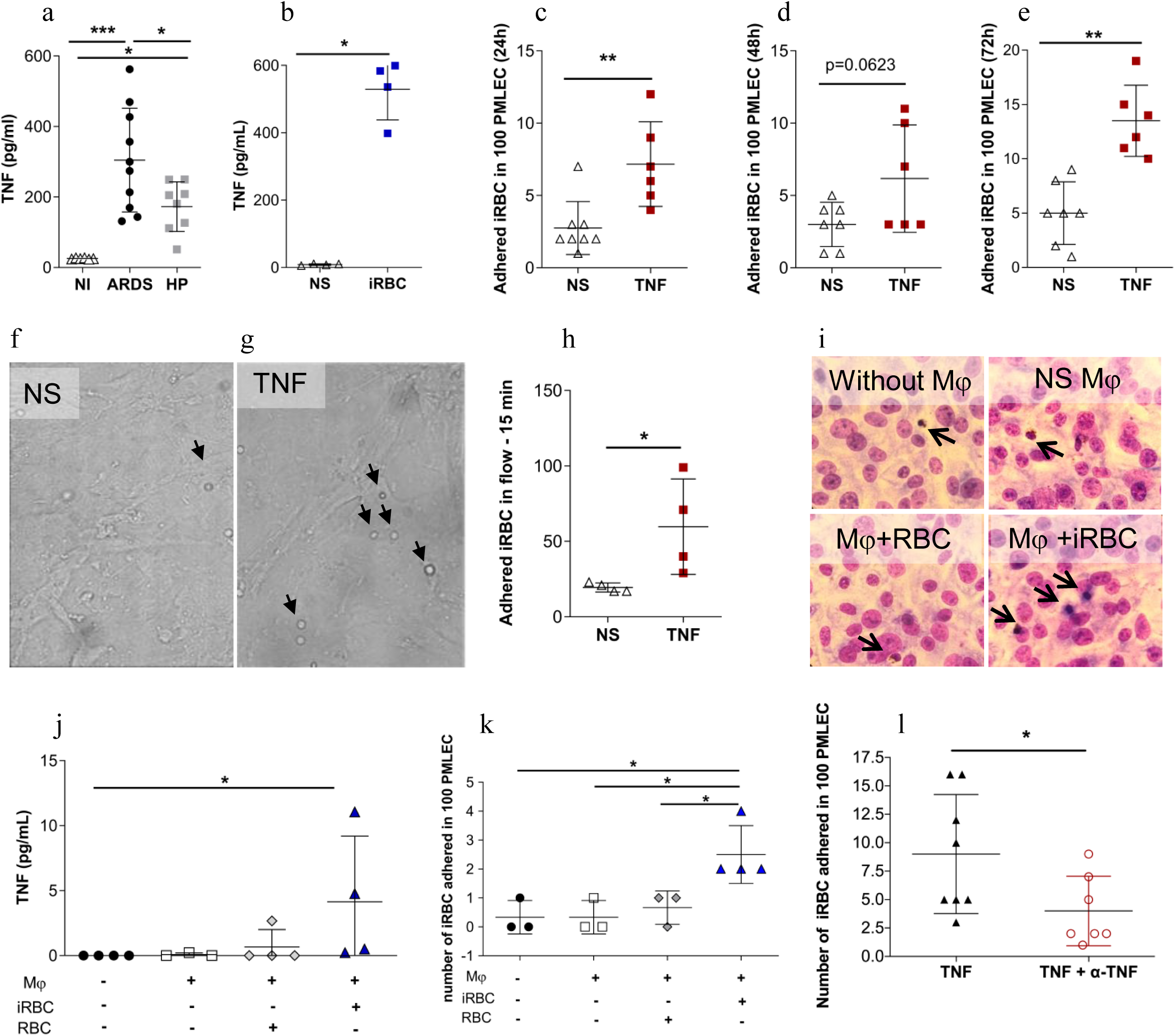
TNF increases *Plasmodium berghei* adherence in the ARDS experimental model. (a) Serum from ARDS- and HP-developing mice was collected on the 7^th^ day postinfection (dpi) and submitted for TNF analysis by ELISA. (b) Primary microvascular lung endothelial cells (PMLECs) from naïve DBA/2 mice were stimulated with infected red blood cells (iRBCs) and TNF was measured in the supernatant. (c-e) PMLECs were also stimulated with TNF (50 ng/ml) and the adhesion of iRBCs to PMLECs under static conditions and (f-h) under flow conditions (after 24 hours of TNF-stimulation) were analysed. (i-k) iRBC adhered in PMLEC in a static condition assay after coincubation with peritoneal macrophages (Mφ), seeded in transwell membranes and coincubated with iRBCs or red blood cells (RBCs) for 24 hours, and then (j) TNF release was measured in the supernatant. (k) PMLECs (that had contact with the Mφ supernatant) were incubated with iRBCs having their adhesion analyzed. (l) iRBC adherence was assessed in TNF-stimulated cells and TNF-neutralized antibodies plus TNF. Bars represent the average ± SEM, Graphics a: one-way ANOVA from two grouped experiments; (b-h) Mann-Whitney test representative of (b and h) two independent experiments and (c-e and l) two grouped experiments; (j and k) Kruskal-Wallis test representative of two independent experiments; Bars represent the average ± SD (*p<0.05; **p<0.01; ***p<0.001). ARDS: acute respiratory distress syndrome; HP: hyperparasitemia; Arrows in f, g and i indicates iRBC adhered in PMLEC.

### EPCR could contribute to *P. berghei* cytoadherence

Looking for adhesion molecules that could be involved in PbA-iRBC cytoadherence in ARDS pathogenesis, we observed that ARDS-developing mice showed higher levels of VCAM and ICAM-1 in lungs than HP-developing or noninfected mice (NI) as analyzed by immunohistochemistry (Figure S2 a-c). Additionally, recombinant TNF upregulated the mRNA expression of ICAM-1 and VCAM in PMLECs (Figure S2 d-i). However, the most interesting finding was the upregulation of mRNA EPCR expression in the lungs of ARDS-developing mice compared to NI mice (28.48-fold increase) and HP-developing mice (13.16-fold increase) (Figure 3a). There was also an increase in soluble EPCR (sEPCR) (Figure 3b) in serum and EPCR protein in lungs (Figure 3c) in ARDS-developing mice when compared to HP-developing mice. We further investigated the influence of TNF on the regulation of EPCR expression in PMLECs. TNF-stimulated cells showed an upregulation of EPCR expression at 48 and 72 hours, but this was not seen with PbA-iRBC stimulation (Figure 3 d-f). Moreover, EPCR knockdown with siRNA transfection (siEPCR) in PMLECs reduced EPCR expression compared to that in nontransfected PMLECs (Figure 3g). In addition, in siEPCR cells, iRBC adhesion was reduced compared to TNF-stimulated and nontransfected cells (Figure 3h).

**Figure 3:**
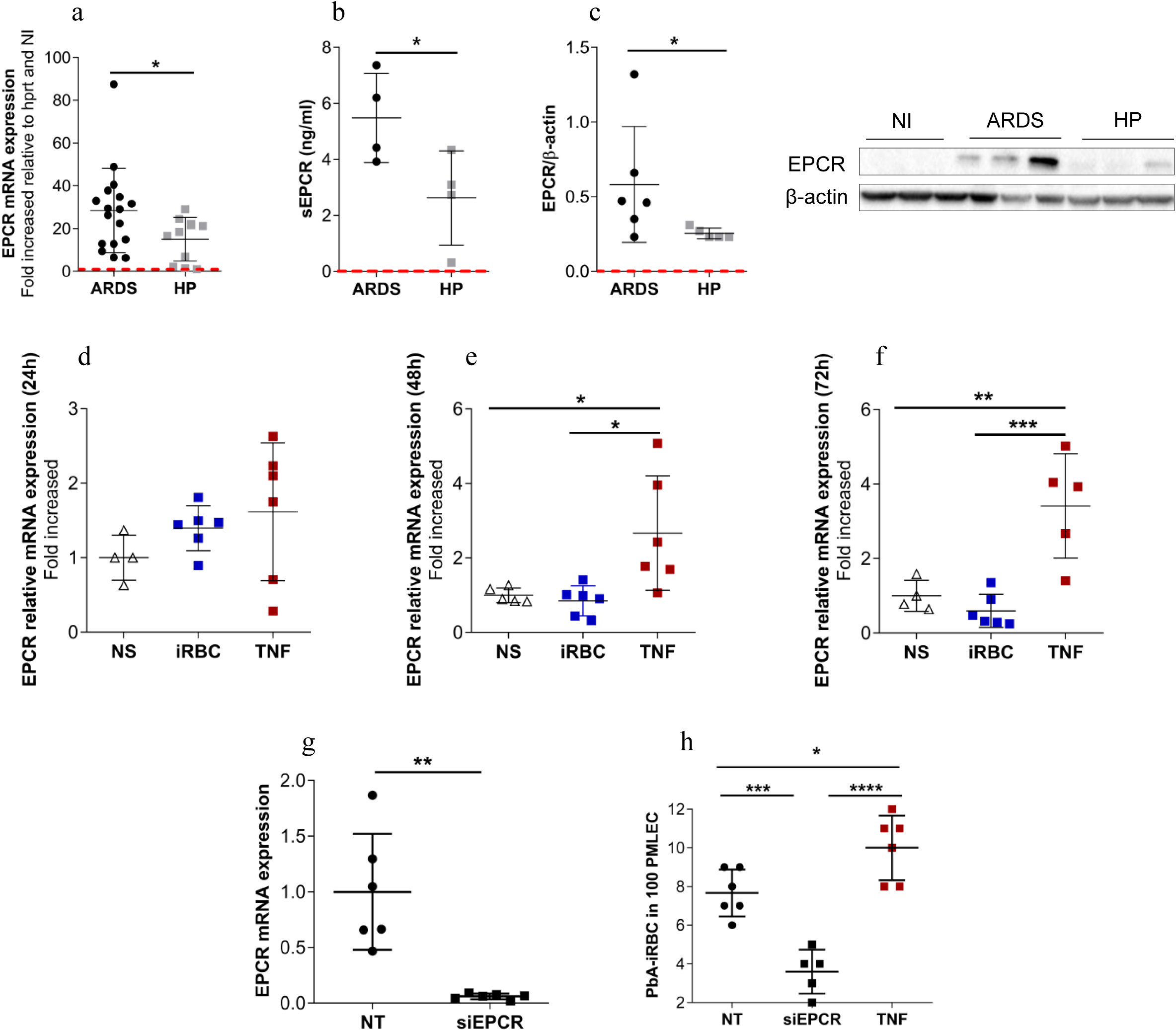
EPCR could contribute to *P. berghei* cytoadherence. Lungs and serum from ARDS- and HP-developing mice were analyzed by (a) EPCR mRNA expression, (b) soluble EPCR (sEPCR) in serum and (c) EPCR protein in lungs, on the 7^th^ day postinfection. (d-f) Primary microvascular endothelial cells (PMLECs) were stimulated with either infected red blood cells (iRBCs) or TNF (50 μg/ml) for (d) 24, (e) 48 and (f) 72 hours to analyze EPCR mRNA expression. (g) PMLECs were subjected to EPCR knockdown with siRNA, and (h) iRBC adherence in PMLECs was evaluated. (a, b and d) Unpaired t-test representative of (a) three grouped, (b) two independent and (d) two grouped experiments; (c) Mann-Whitney test representative of two independent experiments; (e-h) One-way ANOVA from two grouped experiments. Bars represent the average ± SD (*p<0.05; **p<0.01; ***p<0.001; ****p<0.0001; n=4-7 replicates/group). ARDS: acute respiratory distress syndrome; HP: hyperparasitemia; NS: nonstimulated cells; NT: nontransfected cells. Red dashed lines: noninfected mice.

### Dexamethasone reduces VEGF, TNF and EPCR, protecting mouse lungs and PMLECs from increased vascular permeability

Once TNF seemed to be important to induce PbA-iRBC adherence and consequently ARDS development, DBA/2 mice were infected with PbA-iRBC and treated with dexamethasone (80 mg/kg) on days 5 and 6 post-infection to investigate whether it could protect them from lung injury.

Dexamethasone treatment decreased TNF concentrations in the plasma (Figure 4a) apart from IL-6 and IL-33 concentrations in treated mice compared to those of nontreated mice (Figure S3 a and b). Dexamethasone also downregulated EPCR (Figure 4b) and VCAM (Figure S2 c) expression but not ICAM expression (Figure S2 d) in the lungs of infected mice. The soluble EPCR concentrations in the serum and bronchoalveolar lavage (BAL) are shown (Figure 4c and d). The concentration of VEGF, an essential factor to increase vascular permeability and ARDS development, also decreased after dexamethasone treatment in the BAL of infected mice on the 7^th^ dpi (Figure 4e).

**Figure 4:**
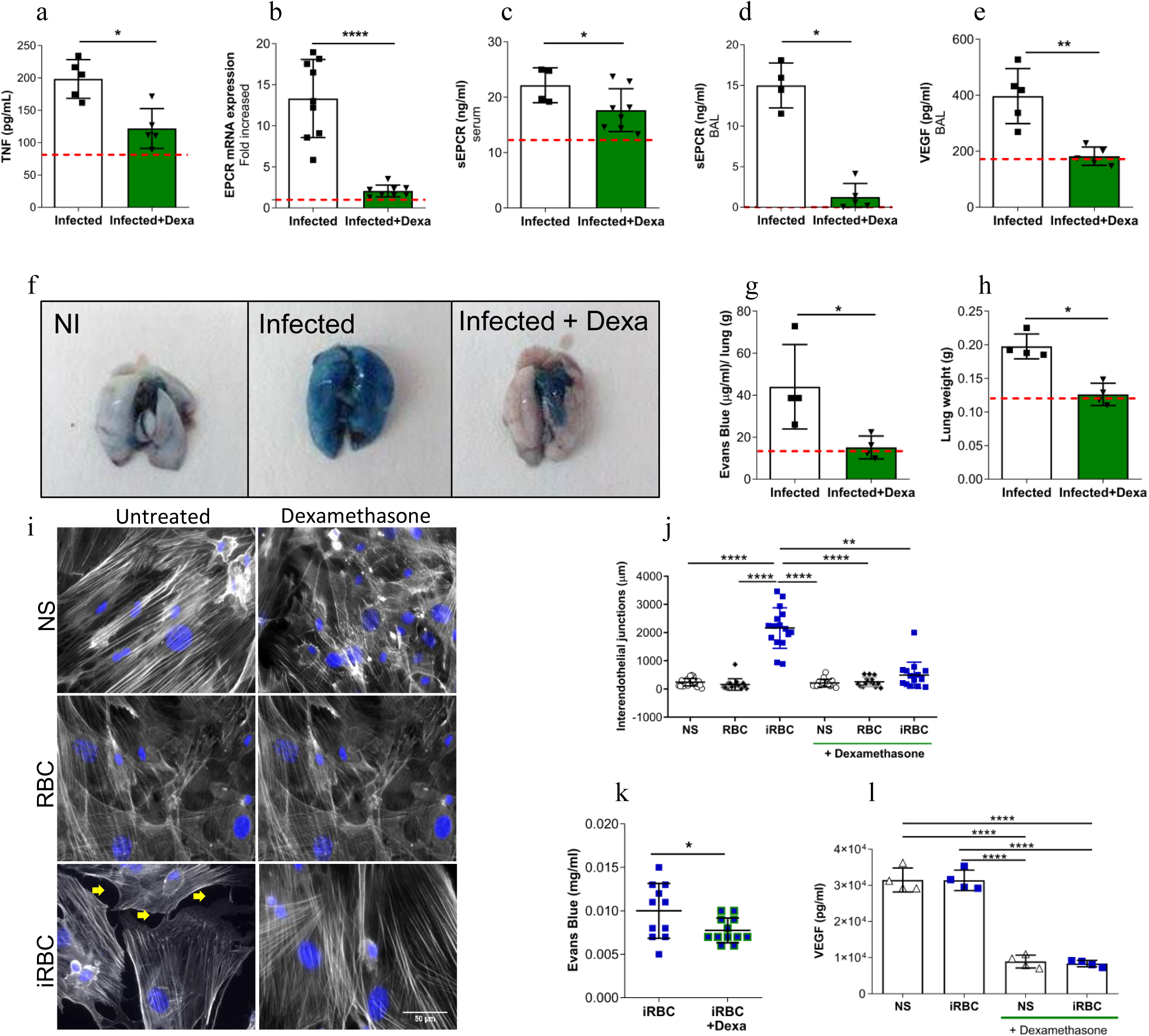
Dexamethasone reduces VEGF, TNF and EPCR, protecting mouse lungs and PMLECs from increased vascular permeability. (a-h) *Plasmodium berghei* ANKA-infected DBA/2 mice were treated with dexamethasone and their serum and perfused lungs were analyzed, on the 7^th^ day postinfection. (a) The TNF concentration in the serum, (b) EPCR mRNA expression in the lungs, (c) sEPCR concentration in both sera and (d) bronchoalveolar lavage (BAL) and (e) VEGF concentration in BAL were evaluated. (f-g) Evans Blue (EB) accumulation was measured in the lungs and (h) weight lungs was analyzed after perfusion. (i-j) Primary microvascular lung endothelial cells (PMLECs) were treated, or not, with dexamethasone and, subsequently, coincubated with either infected red blood cells (iRBCs) or red blood cells (RBCs). (i) PMLECs were stained for actin microfilaments (gray), and nuclei were stained with Hoechst stain (blue), and (j) openings in the interendothelial junctions (OIJ) (indicated by yellow arrows) were compared among all groups. (k) The permeability of dexamethasone-treated PMLECs was related using Evans blue concentration in a Transwell assay. (l) VEGF levels were measured in these groups. Infected+Dexa: *P. berghei* ANKA-infected and dexamethasone-treated mice. Graphics (a, c, d, g and h) Mann-Whitney test representative of (a, d, g and h) two independent and (c) two grouped experiments. (b, e and k) Unpaired t-test from (b) two grouped, (e) from two independent and (k) three grouped experiments; (i) Kruskall-Wallis test from three grouped experiments; (l) One-way ANOVA representative of two independent experiments. Bars represent the average ± SD (*p<0.05; **p<0.01; ****p<0.0001). Red dashed lines: noninfected mice.

Consequently, we analyzed lung vascular permeability, and dexamethasone-treated infected mice were protected compared to nontreated mice (Figure 4f-h). To clarify the mechanism that reduced lung permeability, the openings of interendothelial junctions in PMLECs were evaluated through actin analysis. The PMLECs stimulated with iRBCs increased the spaces between the interendothelial junctions and led to disruption and reorganization in the actin filaments, which were protected by treatment with dexamethasone (Figure 4i and j). iRBC-stimulated PMLECs also demonstrated more permeability for Evans Blue dye in a Transwell assay compared to dexamethasone-treated PMLECs (Figure 4k). Additionally, dexamethasone-treated PMLECs showed a reduction in VEGF concentration compared to cells untreated with dexamethasone (Figure 4l). With everything considered, these data suggest that dexamethasone is capable of reducing inflammation and decreasing EPCR levels and vascular permeability in PMLECs and the lungs of DBA/2 mice.

### Anti-inflammatory drug protects mice from ARDS but not from malaria infection

Dexamethasone-treated mice improved their respiratory parameters [respiratory frequency, tidal volume and enhanced pause (Penh)] compared to infected nontreated mice on the 7^th^ dpi (Figure 5 a-c). On the other hand, parasitemia levels increased in the peripheral blood of dexamethasone-treated mice on the 7^th^ dpi (Figure 5d), and these mice succumbed between 8 and 20 dpi (Figure 5e) with characteristics of HP without edema and lower inflammatory infiltration according to the histopathological analyses (Figure 5 f-g) and count of iRBC data (Figure 5d).

**Figure 5:**
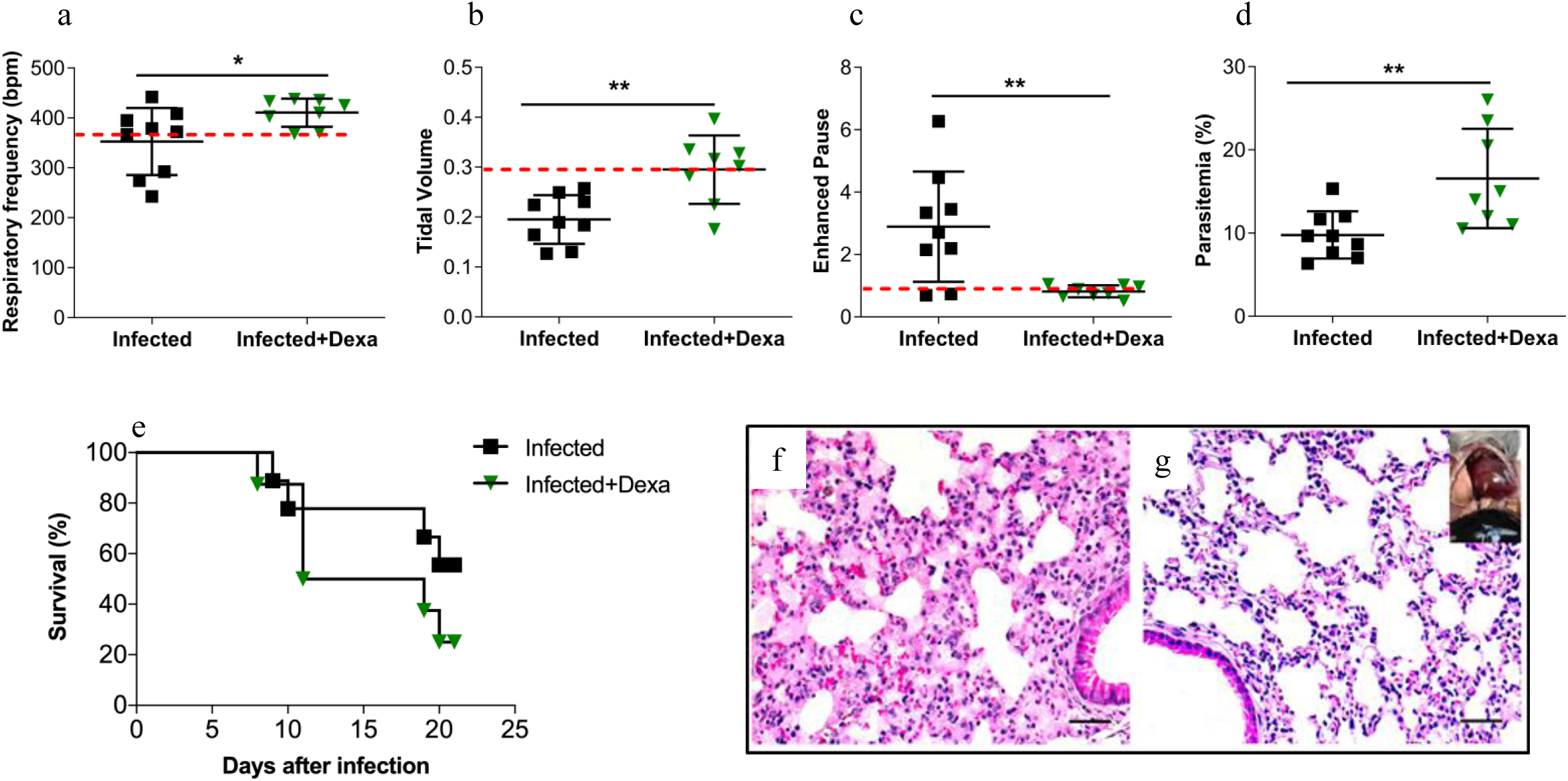
Dexamethasone protects mice from ARDS but not from malaria infection. *Plasmodium berghei* ANKA-infected DBA/2 mice were treated with dexamethasone and compared to infected-untreated mice, on the 7^th^ day post-infection for respiratory parameters: (a) respiratory frequency, (b) tidal volume and (c) enhanced pause (Penh) analyzed in whole body plethysmography chambers. (d) Parasitemia (%) in the peripheral blood and (e) the survival curves were compared between groups. Representative image of histological analysis of lungs from (f) an infected mouse and (g) an Infected+Dexa mouse (Scale bar: 50 µm). Infected+Dexa: *P. berghei* ANKA-infected and dexamethasone-treated mice. Graphics (a-d) Unpaired t test, representative data from two independent experiments. Bars represent the average ± SD. (*p<0.05; ** p<0.01). Red dashed lines: noninfected mice.

## Discussion

In the present study, we describe the cytoadherence of PbA-iRBCs in microvascular lung endothelial cells in ARDS-developing mice, as well as potential contribution of the EPCR in this mechanism and, consequently, in ARDS pathogenesis. An important advantage of this study is that our animal model reproduces the physiopathological aspects found in human ARDS patients [12,13,32].

We have previously shown that PbA-infected DBA/2 mice can develop ARDS (between 7 to 12 days after infection) or develop HP later. ARDS-developing mice show, from 7^th^ dpi, interstitial and lung edema, higher vascular permeability, lung opacification, worse breathing capacity unlike HP-developing mice [12]. ARDS-developing mice also produce more neutrophils-attracting chemokines, myeloperoxidase and reactive oxygen species resulting in increase of neutrophils in lungs and BAL [14]. In this work, we investigated how PbA-iRBC interaction with endothelial cells could mediate some of those changes in the DBA/2 model, leading to one of the two different phenotypes.

It is well known that mature forms of *P. falciparum*-iRBCs can adhere to the microvasculature of different organs, which is an important feature in the pathogenesis of severe malaria in humans [5,33]. However, there are only a few studies focusing on cytoadherence in ARDS [11,29,34,35], and the mechanism of this is not entirely known. Previously, we found that ARDS-developing mice showed a higher level of parasites in their lungs than HP-developing mice, which may be important to induce malaria [14]. Our work reinforces that PbAluc-iRBC accumulated in the lungs of ARDS-developing mice, which could contribute to pathogenesis. Additionally, ARDS-developing mice showed a larger hemozoin (Hz) area in the lungs than HP-developing mice on the 7^th^ dpi, which is consistent with the onset of ARDS development as patients from Thailand with ARDS development presented a significant amount of pulmonary Hz [36]. Some previous works suggest that Hz, which is released from the food vacuole into circulation during erythrocyte lysis, is rapidly taken up by circulating monocytes and tissue macrophages, inducing the production of pro-inflammatory mediators [37,38]. In addition, Hz linked by *Plasmodium* DNA is a ligand for TLR 9, which activates innate immune responses *in vivo* and *in vitro*, resulting in the production of cytokines and chemokines and the upregulation of costimulatory molecules [39,40] and proinflammatory mediators such as IL-1β via NLP3 inflammasome complex activation [38].

It is known that iRBCs adhere to endothelial cells *in vivo* and *in vitro.* Recently, PbA-iRBCs have been demonstrated to adhere to MVECs (microvascular lung endothelial cells from CBA/Ca mice) *in vitro*, and TNF-stimulated cells expressed more ICAM-1 and VCAM [17]. Other studies showed that endothelial cells stimulated with TNF showed an increase in ICAM-1 expression [34,41], corroborating our results *in vivo* and *in vitro* that recently demonstrated that TNF and ICAM-1 markers were increased in the lungs of patients who had developed the syndrome [36]. The inhibition of *P. falciparum* iRBC adherence in TNF-activated HUVECs incubated with either anti-ELAN-1 mAB BBII or anti-VCAM mAB 4B9 [42] was not observed, suggesting that other molecules could contribute to the adhesion of iRBCs. It is interesting to highlight that the parasites by themselves (PbA-iRBC or PbA lysate) cannot upregulate ICAM-1, VCAM or EPCR expression in DBA/2-PMLECs. In fact, PbA-iRBCs adhere more to TNF-stimulated DBA/2-PMLECs than nonstimulated cells. We previously observed the close contact between PbA-iRBCs and endothelial cell membranes in the lungs of ARDS-developing mice [32]. Nevertheless, the study reported here suggests the mechanism of PbA-iRBC adhesion using DBA/2 mice that had developed ARDS. We found that ARDS-developing mice have approximately 28.5 times more EPCR expression than uninfected mice; they showed more EPCR protein in their lungs than HP-developing mice, and TNF-stimulated PMLECs displayed an upregulation in EPCR expression compared to that of nonstimulated cells (NS) or cells stimulated with iRBCs, indicating that EPCR can contribute to ARDS progression. Similarly, a study indicated that children in Papua New Guinea with severe malaria had higher antibody levels against EPCR binding to CIDRα1 domains compared to uncomplicated malaria and this difference was only found in older children [43]. However, in a recent study, the expression of EPCR decreased in the lungs of *P. falciparum*-infected patients who developed ARDS when compared to those who did not develop ARDS as analyzed by immunohistochemistry. The authors suggest that the change in EPCR together with thrombomodulin and in association with the deposition of hemozoin in the lungs plays an important role in the pathogenesis of ARDS [36]. Additionally, reduced sEPCR serum levels were detected in children with severe malaria, and these levels were higher in cerebral liquid from children with cerebral malaria. In spite of this, sEPCR levels were not related to mortality or neurologic manifestations at discharge or 6-month follow-up [44].

Despite the fact that *P. berghei* ANKA does not express PfEMP1, it can cause severe malaria. PfEMP-homologies proteins are not displayed from *P. berghei* and exported proteins from this parasite species, responsible for iRBC cytoadherence, are not unrevealed yet. On the other hand, *P. berghei*-iRBC have shown close contact with lung endothelial cells in ARDS-developing mice [32]. Nevertheless, a recent malaria proteomic and genetic study shows that *P. berghei* blood stages export diversified collection of proteins [45]. One molecule that have been previously reported to mediate adhesion of iRBC in endothelial cells is the schizont membrane-associated cytoadherence protein (SMAC), that is exported into the cytoplasm of the host erythrocyte. Mutants lacking expression of SMAC show a strongly reduced CD36-mediated iRBC sequestration [46]. Unfortunately, we have not found any correlation between CD36 mRNA levels and ARDS development in our model. Passini *et al*, 2013 identified erythrocyte membrane associated proteins, but related in sequestration [45]. Therefore, further studies should be developed to recognize this *P. berghei*-proteins, especially with potential to connected EPCR adhesion.

With the hypothesis that inflammation is essential to expose adherence molecules and consequently to ARDS pathogenesis, we treated DBA/2 mice with 80 mg/kg of dexamethasone. The 80 mg/kg dose is the therapeutic dose for ARDS treatment in mice observed by Van den Steen in 2010, because lower doses did not protect mice from ARDS and edema [29]. In contrast, dexamethasone was administered to patients suffering from ARDS (different etiologies) at lower doses: 5 mg/day for 4 days in ARDS patients with bacterial pneumonia [47] and 10 mg for 6 hours in ARDS patients with acute leucocyte leukemia [48]. In two well-conducted studies, dexamethasone failed to improve the fatality rate of cerebral malaria [49,50]. Unfortunately, the use of corticosteroids in acute respiratory distress syndrome (ARDS) due to malaria in humans has not been well explored [51]. In our experiments, we observed that dexamethasone-treated mice were protected from ARDS and showed the downregulation in EPCR expression, which indicates that EPCR may be necessary for iRBC adherence and the progression of disease. In other models, using endothelial cells (HUVECs), dexamethasone also had a direct effect on the down-regulation of the EPCR expression [52].

EPCR bind to *Plasmodium* was suggested to block the interaction of EPCR with protein C. [53]. On the other hand, APC bind with EPCR was required to inhibit NETosis [54]. We have previously shown that *P. berghei* and *P.falciparum*-iRBC induced NET formation and its importance in ARDS development [14].

However, there are specific domains of the PfEMP1 subfamily that do not bind to EPCR while others do [20], although the presence of serum inhibits this binding during *in vitro* assays [25]. Nevertheless, recently, a new study showed that using human brain microvascular endothelial cells and 10% human serum did not affect the binding between *P. falciparum*-infected red blood cells and EPCR. In addition, parasites isolated from patients with cerebral malaria displayed higher binding capacity of cytoadherence under flow conditions, compared with isolated from uncomplicated malaria patients [55]. Parasite var transcripts encoding EPCR-binding domains, in association with high load of parasite and low platelet levels, are hardy indicators of cerebral malaria. Patients with cerebral edema showed an increased transcript of parasite PfEMP1 DC8 and EPCR-binding domains. In addition 62B1-1-CIDRa1.7 affected EPCR-APC interaction and the authors proposed a harmful role for EPCR-binding domains in these patients [56]. Therefore, the exact role of EPCR is controversial and needs to be better understood. In addition, dexamethasone-treated mice are less susceptible to increased vascular permeability. Likewise, iRBC-stimulated PMLECs treated with dexamethasone showed less permeability than untreated cells. *P. berghei* NK65-infected C57BL/6 mice treated with the same dose of dexamethasone also showed an increase in peripheral parasitemia and ARDS protection [29] but not an effect on pulmonary vascular permeability.

Based on our results, treatment with dexamethasone, a potent anti-inflammatory drug, decreased the serum concentrations of IL-33, IL-6, and TNF in PbA-infected mice. In malaria, high level of TNF was associated with severe malaria, especially in children. TNF can participate in parasite killing and, consequently, parasitemia reduction. On the other hand, increased TNF levels were observed in children with high-density parasitemia and when parasite clearance occurs, TNF decays. It is described, in a recent review, that TNF polymorphisms could control parasite development in early stages of infection, but can enlarge the probability of patients to develop severe malaria syndromes [57].

TNF is an inflammatory mediator strongly implicated in the development of ARDS [58] signaling through two receptors, p55 and p75, that play differential roles in pulmonary edema formation during ARDS. It was shown by a novel domain antibody (dAb™) that p55 attenuated ventilator-induced lung injury [59] and lung injury and edema formation in models of ARDS induced by acid aspiration [58]. Recently, the selective blockage of TNFR1 (GSK1995057 antibody) inhibited cytokine and neutrophil adhesion molecule expression in activated HMVEC-L monolayers *in vitro* and attenuated inflammation and signs of lung injury in primates [60]. Moreover, treatment with this antibody attenuated pulmonary neutrophilia, inflammatory cytokine release and signs of endothelial injury in BAL and serum samples in healthy humans challenged with a low dose of inhaled endotoxin [60]. However, the precise mechanisms by which TNF is involved in severe malaria remains unknown [57].

IL-33 is a tissue-derived nuclear cytokine from the IL-1 family, and it is highly expressed during homeostasis and inflammation in endothelial, epithelial and fibroblast-like cells, working as an alarm signal released upon cell injury or tissue damage to alert immune cells to express the ST2 receptor (IL-1RL1) [61]. Bronchial IL-33 expression is significantly increased in severe malaria patients with pulmonary edema [62]. Moreover, in PbA-induced experimental cerebral malaria (ECM), IL-33 expression is increased in the brain, and ST2-deficient mice were resistant to PbA-induced neuropathology [63,64]. On the other hand, PbA-infected C57BL/6 mice treated with recombinant IL-33 presented no signs of neurological pathology associated with CM and had reduced production of pro-inflammatory cytokines and chemokines [65].

Circulating IL-6 levels are elevated in nearly all infectious, traumatic, and inflammatory states, including ARDS. Elevated levels of IL-6 are found in the BAL and plasma of patients with ARDS and those at risk [66,67].

Finally, we proposed the essential mechanisms of protection for dexamethasone in experimental malaria (Figure 6), suggesting that PbA-iRBCs induce TNF release by endothelial cells and by Mφ, and then TNF upregulates EPCR expression in PMLECs, increasing iRBC adhesion through the EPCR pathway. The adhesion of iRBCs in pulmonary endothelial cells leads to an increase in gap formation in the interendothelial junction, raising vascular permeability and consequently the formation of edema. Activated alveolar macrophages also produce TNF, which contributes to the activation of endothelial cells and possibly the recruitment of neutrophils [14] and alveolar damage. The drastic reduction in inflammation with dexamethasone treatment may be responsible for the increase in parasitemia and mortality. These data suggest that TNF is essential to increase the expression of EPCR and consequently increase adhesion of iRBCs in malaria-associated ARDS.

**Figure 6:**
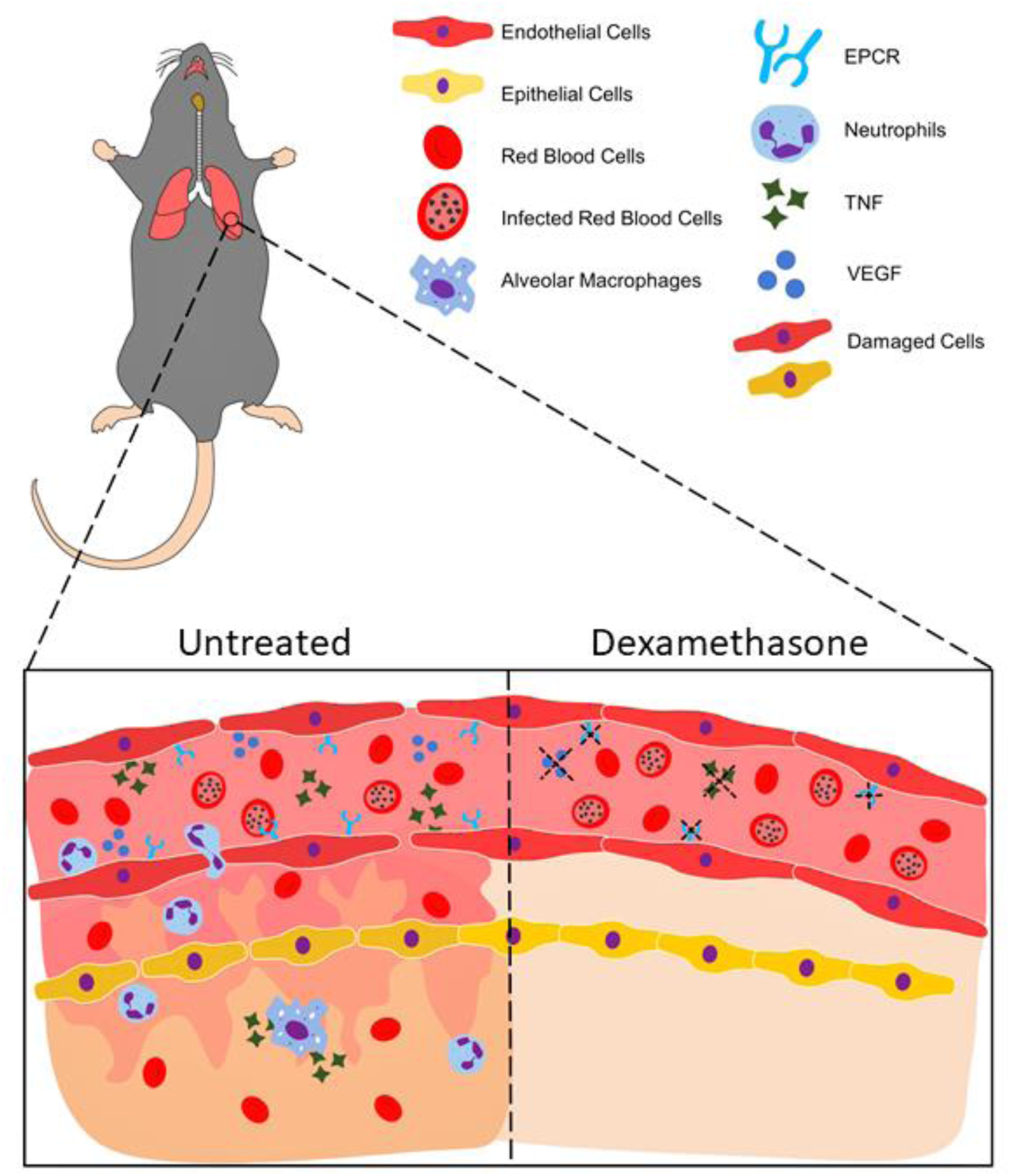
Schematic representation of ARDS development involving adhesion of iRBCs to EPCR. *Plasmodium berghei* ANKA-infected red blood cells (iRBCs) induce TNF release by endothelial and inflammatory cells. TNF upregulates EPCR expression in lung endothelial cells, increasing iRBCs adhesion in these cells through EPCR pathway. The adhesion of iRBCs to lung endothelial cells leads to an increase in gap formation in the interendothelial junction, an increase in vascular permeability with infiltration of inflammatory cells and red blood cells and edema formation. Activated alveolar macrophages also produce TNF, which contributes to the activation of endothelial cells, recruitment of neutrophils and alveolar damage. Treatment with dexamethasone decreases inflammatory cytokine release, including TNF release and, consequently, downregulates EPCR expression in lung endothelial cells; dexamethasone also decreases VEFG released by endothelial cells, protecting mice from gap formations, vascular permeability, inflammatory cell infiltration and alveolar damage.

## Conclusions

Our data together suggest that *P. berghei* infection induces TNF production by inflammatory and endothelial cells, leading to the expression of adhesion molecules such as ICAM-1, VCAM and, especially, EPCR. These results allow us to infer that those factors could contribute to PbA-iRBC cytoadhesion and thus suggest the participation of these mechanisms in the pathogenesis of malaria-associated ARDS. Knowing that inhibition of cytoadhesion mechanisms may improve the prognosis of infection [68], we emphasize that the processes described above may be important in the treatment of ARDS due to *Plasmodium* infection, acting as adjunctive therapy in association with the use of antimalarials, not only making the treatment more efficient but also reducing the morbidity and mortality of these patients.

## Conflicts of Interest

The authors declare that the research was conducted in the absence of any commercial or financial relationships that could be construed as a potential conflict of interest.

## Funding Statement

This work was supported by the São Paulo Research Foundation (Fundação de Amparo à Pesquisa do Estado de São Paulo: FAPESP, Brazil), the Coordination of Improvement of HigherLevel Personnel; (Coordenação de Aperfeiçoamento de Pessoal de Nível Superior: CAPES) and the National Council for Scientific and Technological Development (Conselho Nacional de Desenvolvimento Científico e Tecnológico: CNPq, Brazil). LSO (CAPES and FAPESP 2013/20718-3), MKS (CAPES/PNPD), GCM (FAPESP 2017/00077-4), TCQ (CNPq 131431/2017-0), DD (CNPq 162730/2014-4), DSC (CNPq 133890/2016-3), OM (FAPESP 2013/00981-1), CRFM (FAPESP 2016/07030-3), SE (FAPESP 2014/20451-0 and 2017/05782-8), and CNPQ (455863/2014-8) received fellowship grants.

## Acknowledgments

The authors thank Patricia Mendonça da Silva Amorim, Erika Paula Machado Peixoto, and Bernardo Paulo Albe for their technical assistance.

