## Supplemental figure 1 for "Endothelial Protein C Receptor Could Contribute to Experimental Malaria-Associated Acute Respiratory Distress Syndrome"

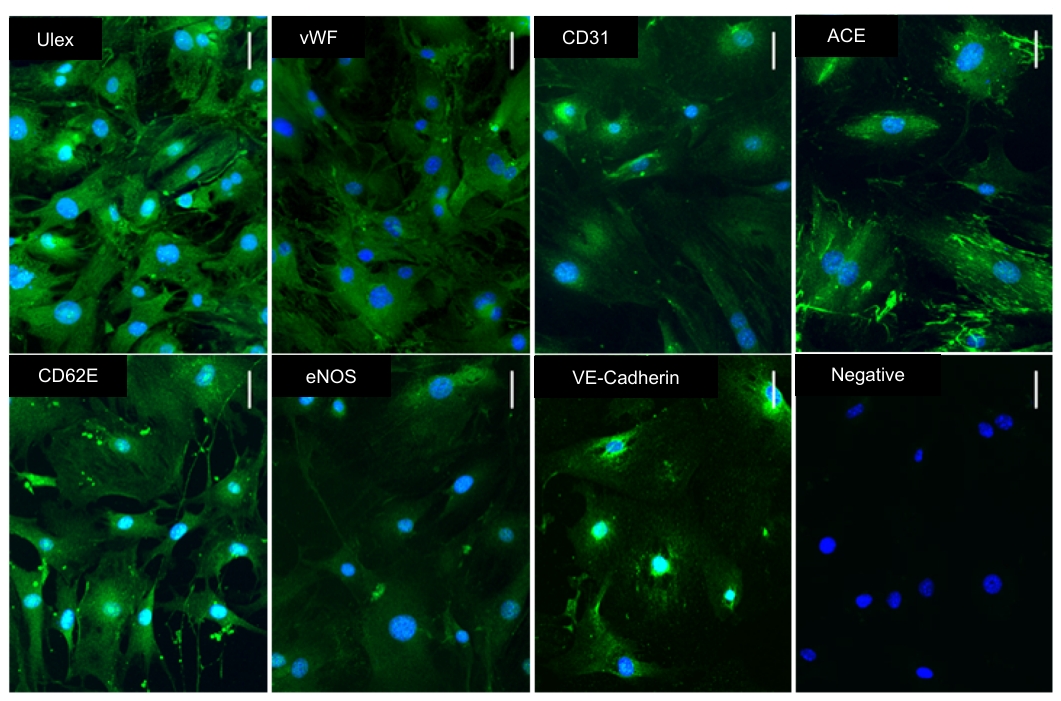

**Supplementary Figure 1:** **Primary microvascular lung endothelial cell characterization.** PMLECs were stained by immunofluorescence with lectin from *Ulex europaeus* and anti-vWF, CD31, ACE, CD62E, eNOS and VE-cadherin antibodies. Nuclei stained with Hoechst stain (blue). PMLECs: primary culture of microvascular lung endothelial cells from DBA/2 mice. (scale bar: 50 µm).
