## Supplemental figure 2 for "Endothelial Protein C Receptor Could Contribute to Experimental Malaria-Associated Acute Respiratory Distress Syndrome"

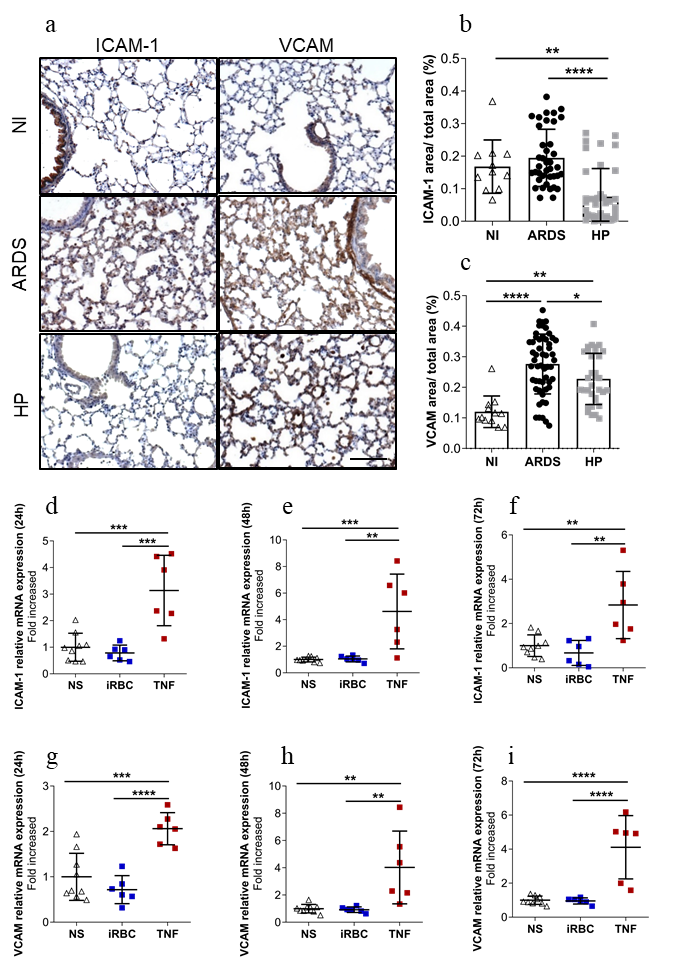

**Supplementary Figure 2:** **ARDS-developing mice show more ICAM-1 and VCAM expression in the lungs than HP-developing or noninfected mice.** (a-c) DBA/2 mice infected with *Plasmodium berghei* ANKA and lung tissues were analyzed on the 7^th^ day post-infection. (a) Representative immunohistochemistry images for ICAM-1 and VCAM in the lungs of NI, ARDS-developing and HP-developing mice (scale bar: 100 µm). (b) Quantification of (b) ICAM-1 and (c) VCAM areas stained by immunohistochemistry. (d-i) Primary microvascular lung endothelial cells from uninfected DBA/2 mice (PMLECs) were stimulated with *Plasmodium berghei* ANKA-infected red blood cells (iRBCs) or with recombinant TNF and analyzed for (d-f) ICAM-1 mRNA and (g-i) VCAM mRNA by qRT-PCR. Bars represent the average ± SD. (b-i) One-way ANOVA from two grouped experiments. (*p<0.05; ** p<0.01; *** p<0.001). ARDS: acute respiratory distress syndrome; HP: hyperparasitemia; NI: non-infected mice; NS: non-stimulated cells.
