## Supplemental figure 3 for "Endothelial Protein C Receptor Could Contribute to Experimental Malaria-Associated Acute Respiratory Distress Syndrome"

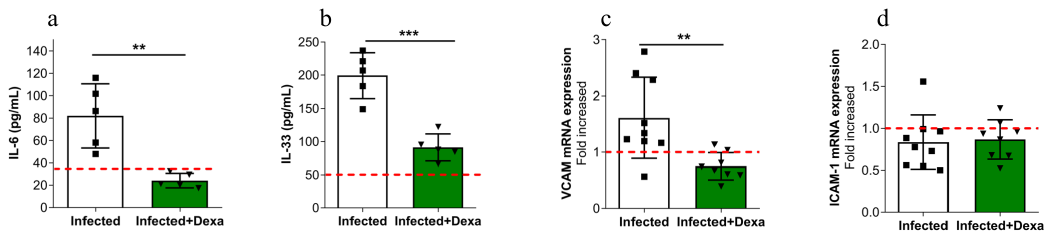

**Supplementary Figure 3:** **Dexamethasone reduced inflammation and VCAM expression but did not decrease ICAM-1 expression in *P. berghei*-infected mice.** *Plasmodium berghei* ANKA-infected DBA/2 mice were treated with dexamethasone and compared to infected-untreated mice. Serum and lung tissues were analyzed on the 7^th^ day post-infection. Serum levels of (a) IL-6 and (b) IL-33 analyzed by ELISA. mRNA expression from lung tissues was evaluated for (c) VCAM and (d) ICAM-1 by qRT-PCR. Bars represent the average ± SD (** p<0.01; *** p<0.001). (a-b) Unpaired t test representative of two independent experiments; (c-d) Mann-Whitney test from two grouped experiments. Infected+Dexa: *P. berghei* ANKA-infected mice and treated with dexamethasone. Red dashed lines: noninfected mice.
